# Application of clinical genomic sequencing among Chinese advanced cancer patients to guide precision medicine decisions

**DOI:** 10.1101/367854

**Authors:** Shunchang Jiao, Yuxian Bai, Chun Dai, Xiaoman Xu, Xin Cai, Guan Wang, Jinwang Wei, Bing Wu, Wending Sun, Qiang Xu

**Author notes:** **Correspondence to** Qiang Xu GenomiCare Biotechnology (Shanghai) Co., Ltd., 5th Floor, Building 2, No. 111 Xiangke Road, Shanghai, China 201210. Shunchang Jiao and Yuxian Bai contribute equally to this work. We state that this article has been submitted to Personalized Medicine.

## Abstract

**Purpose:** A number of studies have suggested that high-throughput genomic analyses might improve the outcomes of cancer patients. However, whether integrative information about genomic sequencing and related clinical interpretation may benefit Chinese cancer patients with stage IV disease to date has not investigated.

**Methods:** Targeted gene panel and whole exome of tumor/blood samples in > 1,000 Chinese cancer patients were sequenced. Then we provided patients and their oncologists with the sequencing results and a clinical recommendation roadmap based on evidence-based medicine, defined as CWES. Only patients with stage IV disease who failed the previous treatment upon receiving the CWES reports were included for analyzing the impact of CWES on clinical outcomes in 1-year follow-ups.

**Results:** We identified the mutational signatures of 953 Chinese cancer patients, with some being unique. Approximately 88.6% of patients had clinically actionable somatic genomic alterations. We successfully followed up 22 stage IV patients. Of these, 11 patients treatment followed the CWES reports defined as group A. Eleven patients received the next treatment, but did not follow the CWES suggestions, and are defined as group B. The types of therapies before CWES were similar in the two groups. The median PFS of group A was 12 months and 45% patients failed this round of therapy. The median PFS of group B was 4 months and 91% of patients failed the treatment.

**Conclusion:** The current study suggested that CWES has the potential to help explore the clinical benefits in multiple line therapies among advanced stage tumor patients.

## Introduction

With the development of targeted and immunotherapy, the era of precision medicine is upon us. Precision medicine, sometimes referred to as “personalized medicine”, can tailor disease prevention and medical treatment according to differences in the patients’ genomics, habits, customs and environments. These treatments allow cancer patients to live longer and achieve a better lifestyle. For example, the emergence of EGFR-targeted drugs has greatly improved the survival of lung cancer patients. Compared to carboplatin plus paclitaxel, gefitinib treatment increased the objective response rates (ORRs) from 47.3% to 72.1%, and progression-free survival (PFS) from 6.3 to 9.5 months. In the EURTAC trial, erlotinib was associated with a significant improvement in ORRs (58% vs 15%) and PFS (9.7 vs 5.2 months) compared to platinum doublet therapy (Fukuoka et al. 2011; Rosell et al. 2012; Thongprasert 2009; Zhou et al. 2011). Targeted therapies and immunotherapy greatly improved the survival and quality of life of these cancer patients.

Targeted therapies and immunotherapy are aimed at specific patients. Until January 23, 2018, the FDA approved nearly 40 companion diagnostic devices for 26 different drugs applications (Silva et al. 2018). These companion diagnostic devices are single gene tests or panel tests. Carcinogenesis is complex, including the multistep process of genetic mutations and tumor heterogeneity. These companion diagnostic devices are only available at present for a limited number of genes. Various clinical studies have reported that the analysis of comprehensive characterization of genome changes produced significant clinical benefits for cancer patients (Takeda et al. 2015; Tsimberidou et al. 2014). In addition, our understanding of cancer has been updated very rapidly. For example, clinical studies have found patients with MDM2 family amplification or EGFR aberrations that became hyperprogressors after immunotherapy (Kato et al. 2017). It is not easy to keep valuable information constantly updated for limited tumor specimens using conventional methods when analyzing tumor specimens.

Although the exome indicates the protein encoding region of the human genome and represents < 2% of the human genome, it contains ∼85% of the known pathogenic variants (van Dijk et al. 2014). Therefore, whole exome sequencing (WES) can provide comprehensive information about the diagnosis, treatment, and prognosis of individual cancer patients. Furthermore, WES can provide a much higher coverage to solve the problem of high-level heterogeneity and mixing with normal tissues in obtained tumor specimens. The use of WES has reduced the sequencing cost (Clark et al. 2011; Roychowdhury and Chinnaiyan 2016). Currently, many clinically-oriented analysis companies have reduced the cost of WES to a price that patients and clinicians can accept. But it spends a bit longer than the panel sequencing. Therefore, the gene sequencing of this study combined targeted gene panel with high coverage level and WES technology.

Of 154 FDA approved cancer drugs, they comprise 17% cytotoxic agents, 25% broadly cytotoxic agents inhibiting, at least partially, protein molecules and 55% that target clear mechanistic protein molecules. The third class of drugs are mostly targeted and immunotherapy drugs (Santos et al. 2017). Except for FDA approved drugs, there are many available and investigational therapies for cancer. These data show explosive growth. Timely accession to increasing treatment information by oncologists and patients will be clearly beneficial for individualized treatment regimens.

In the present study, we have provided targeted gene panel sequencing, WES results and clinical recommendation roadmaps based on evidence-based medicine, also called CWES, for more than 1,000 Chinese cancer patients and their physicians. We identified the mutational signatures of 953 Chinese cancer patients and compared with the data of TCGA. Then, we conducted a follow-up survey in order to assess the clinical benefits of CWES.

## Methods

### Cohort of patients

Between October 2015 and March 2016, we obtained over 1,000 cancer samples from more than 70 hospitals across 20 provinces in China, including 655 formalin-fixed, paraffin-embedded (FFPE) and 524 fresh tumor tissues, making a total of 1,179 samples tested. Matched blood samples were collected as controls. Due to unqualified samples, failure detection and for other reasons, a total of 953 cases were successfully completed CWES analysis.

Only patients with stage IV disease who failed the previous treatment upon receiving the CWES reports were included in the analysis for the clinical benefits of CWES. About 50% tumor specimens were IV, but most of these were only samples, not patients that can be included in the study. In addition, patients who were unable to contact, died, and were unwilling to participate in the study were excluded. We followed 25 patients. Follow-up was conducted every 3 months after patients and oncologists received the CWES reports. Follow-up started from the first half of 2016 until June 2017. Of 13 patients who accepted treatment followed the CWES reports, 1 patient stopped treatment due to adverse drug reactions, and 2 patients were lost to the study. A total of 12 patients accepted treatment without following the CWES reports, and 1 patient was lost.

We obtained written informed consent from the donors for the use of genomic and clinical data for research purposes. All procedures performed in studies involving human participants were in accordance with the ethical standards of the institutional and/or national research committee and with the 1964 Helsinki declaration and its later amendments or comparable ethical standards.

### Panel sequencing, DNA library preparation and whole exome sequencing, and panel and whole exome mapping

Targeted gene panel is a custom multiplexed PCR amplification-capture based assay, including BRAF, EGFR, ERBB2, KIT, KRAS, MET, NRAS, PIK3CA, ALK, RET and ROS1 which are druggable by China food and drug administration (CFDA) approved for treatment. The whole exome contains exonic sequences of 27,000 genes. These genes and whole exome of tumor samples and matched blood samples were sequenced. For the panel, preparation of amplicon libraries was performed by Ion AmpliSeq On-Demand Panels, then were sequenced by Ion Torrent ™ Ion S5 XL Systems. Panel sample coverage level was 4000X. For the whole exome, DNA was fragmented and hybridized to Agilent SureSelect Human All Exome kit V5. The Illumina Xten platform sequenced exome shotgun libraries, generating pair-end reads with sizes of 150 * 2 bp. The average sequence coverage of the whole exome was 200X. Illumina CAVSAVR completed the image analysis and base calling, using default parameters. Then reads with sequences matching the sequencing adaptors and low-quality reads with matches were deleted and high-quality reads obtained. These reads were aligned to the NCBI human reference genome hg19 using the Burrows-Wheeler Aligner alignment algorithm. In addition to WES, we also used panel detection. First, we can quickly obtain information on important driver genes; second, coverage level of 4000X can detect mutations with low mutation frequency.

### Analysis of panel and WES data

We used the Genome Analysis Toolkit (GATK) (version 3.5) to process reads. Localized insertion-deletion indel realignments were performed by GATK. GATK Realigner Target Creator identified regions that needed to be realigned.

For SNV calling, the MuTect algorithm was applied to identify candidate somatic single-nucleotide variants in tumors compared with a matched control blood sample from 1 patient. SNV annotation was performed by ANNOVAR. We used dbNSFP31 to predict nonsynonymous mutations of the encoded proteins.

For indel detection, tumor and blood samples were analyzed with Varscan Indel. Candidate somatic indels were identified if they satisfied the following conditions: they were supported by at least 5 reads; the number of supporting reads divided by the maximum of the read depth at the left and right breakpoint positions was larger than 0.05; the Integrative Genomics Viewer manually checked all somatic indel calls.

For CNV detection, CNVs were generated referring to this method (Sathirapongsasuti et al. 2011). We set all parameters to the default setting for filtering samples, and provided the data for normalization via XHMM. SVD was set to 1 and the bin size was set to 50–60, according to the average coverage depth for CNVnator.

All procedures follow the Molecular Pathology Clinical Practice Guidelines and Reports (Li et al. 2017).

### Tumor mutational burden (TMB)

TMB was defined by the total number of non-synonymous somatic mutations. The determination of TMB followed this method (Chalmers et al. 2017). We defined tertile of TMB of each cancer specie as threshold of high TMB according to prior study (Peters et al. 2017).

### Microsatellite instability (MSI)

All autosomal microsatellite tracts containing 1-5 bp repeating subunits in length and comprising 5 or more repeats in GRCh37/hg19 were identified using MISA (http://pgrc.ipk-gatersleben.de/misa/misa.html). The detailed methodology was followed according to reference (Hause et al. 2016). ≥3.5% of unstable microsatellite sites was defined high MSI according to prior report (Niu et al. 2013).

### Clinical relevant treatment recommendations

We provided clinical recommendation roadmaps for cancer patients and their physicians based on the following data:

1. CFDA and FDA approved drugs for matched tumor types.
2. Professional guidelines as therapy recommended drugs for specific types of tumors.
3. Drugs were available as off-label treatment for the specific molecular alteration in a non-approved tumor type.
4. Investigational clinical trials provided agents based on an identified molecular alteration.

## Statistical analysis

The primary objective was to evaluate whether patients could benefit from following the CWES reports. The primary endpoint was PFS that was defined as the time from the beginning of next round treatment after the patient received the CWES reports to disease progression or death resulting from any cause. Tumor response and progression were assessed by physician notation who accorded to RECIST 1.1 (Sathirapongsasuti et al. 2011). PFS were estimated and plotted by the method of Kaplan and Meier (log-rank). The chi-squared (χ^2^) test or Fisher’s exact test is used with categorical data. P value less than 0.05 was statistically significant. Statistical tests were analyzed using SPSS software (Version 23.0.0).

## Results

### Description of the sequencing cohort

We obtained 1,197 tumor samples, and successfully completed 953 CWES reports (*Figure 1*). These tumors encompassed 4 principal tumor types and more than 18 detailed tumor types. The 4 principal tumor types were colon cancer (27%), lung cancer (26%), breast cancer (16%), and gastric cancer (9%), which accounted for about 78% of the total, covering the main types of cancers found in Chinese patients (Chen et al. 2016). The next most common types of cancer were renal cell carcinoma (4%), hepatocellular carcinoma (4%) and sarcoma of soft tissue (4%). In addition, compared to lung squamous cell carcinoma (LUSC) andother types of lung cancer, we detected a large degree of lung adenocarcinoma (LUAD), which is in line with the distribution of lung cancer in China (Chen et al. 2016). In breast cancer, the number of patients with HER2 positive, HER2 negative and HER2 unknown was similar (*Figure 2*).

**Figure 1.**
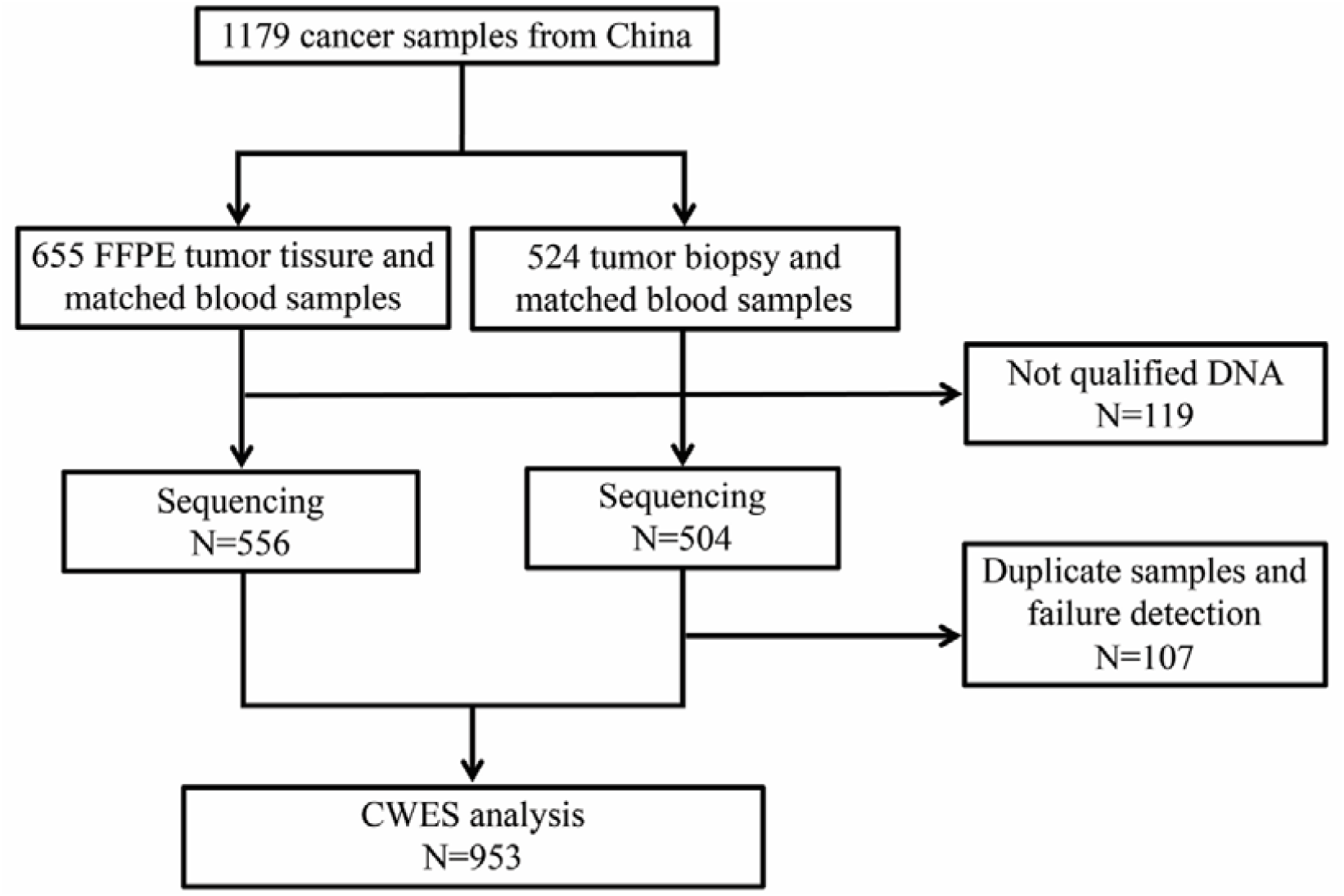
Flowchart of the experimental protocol.

**Figure 2.**
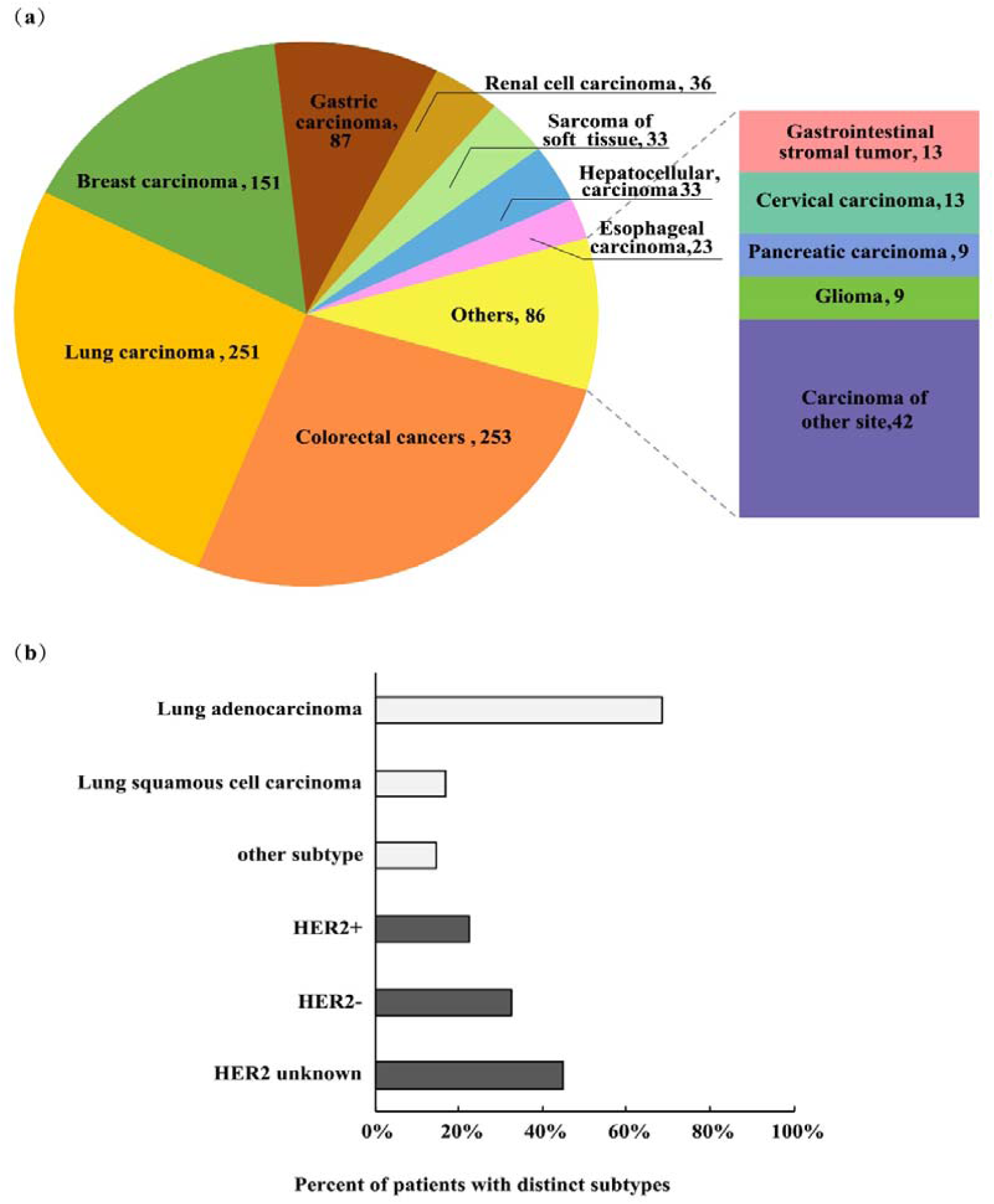
Overview of the Chinese patients cohort. (a) Distribution of tumor types from 953 Chinese cancer patients. Cases included 4 principal tumor types and more than 18 detailed tumor types. (b) Percent of Chinese patients with distinct subtypes of lung and breast cancer. In lung cancer, lung adenocarcinoma is the main subtype, including 171 cases (67%). The distribution of HER2^+^, HER2^-^ and HER2 unknown is similar in breast cancer.

### Mutational signatures of Chinese cancer patients

First, we analyzed the main somatic mutations and CNV of Chinese cancer patients. Some were consistent with the TCGA data, but others were unique. APC and KRAS are the main driving mutations of colorectal cancer in Chinese and foreign populations, but the mutation rate is slightly lower than that of TCGA. Except for TP53, there are no high-frequency mutations in gastric cancer in the Chinese population. CDH1 is the second high-frequency somatic mutation gene in China, the mutation frequency of CDH1 is similar to that of TCGA (9.63% vs 10%), but CDH1 is not the main driving gene in TCGA. In LUSC, the most common changes in CNV were amplification of PIK3CA and loss of CDKN2A, with little difference in the ratio between Chinese and foreign populations. In LUAD, we found that the EGFR and KRAS mutation rates were significantly different between China and TCGA. The frequency of EGFR mutations was 45.97%, which is more than 3 times that of TCGA (14.35%), and the mutation frequency of KRAS was one-third that of TCGA (11.41% % vs 32.61%) (*Figure 3*). These data indicate that the main pathogenesis of Chinese LUAD is different from foreigners.

**Figure 3.**
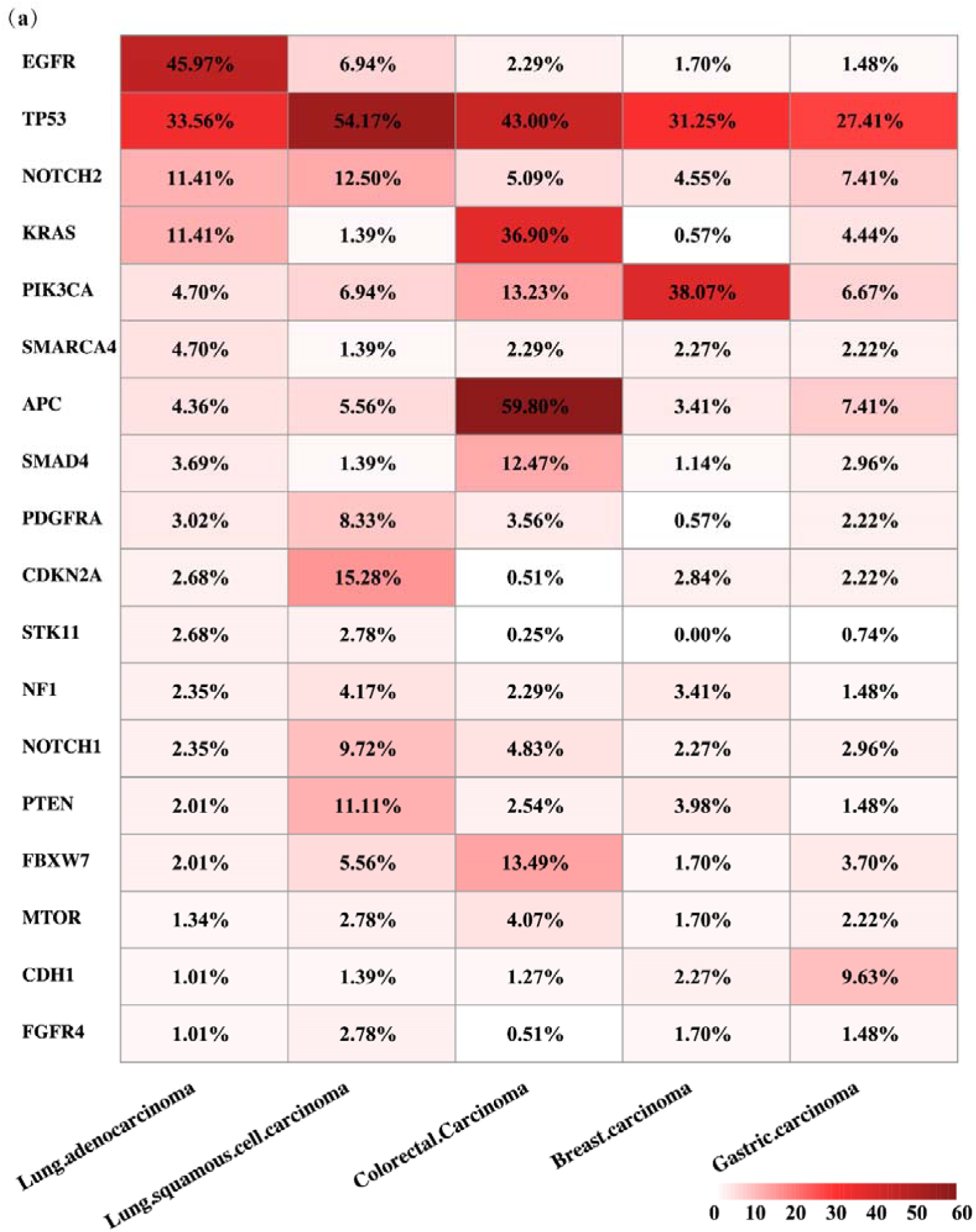
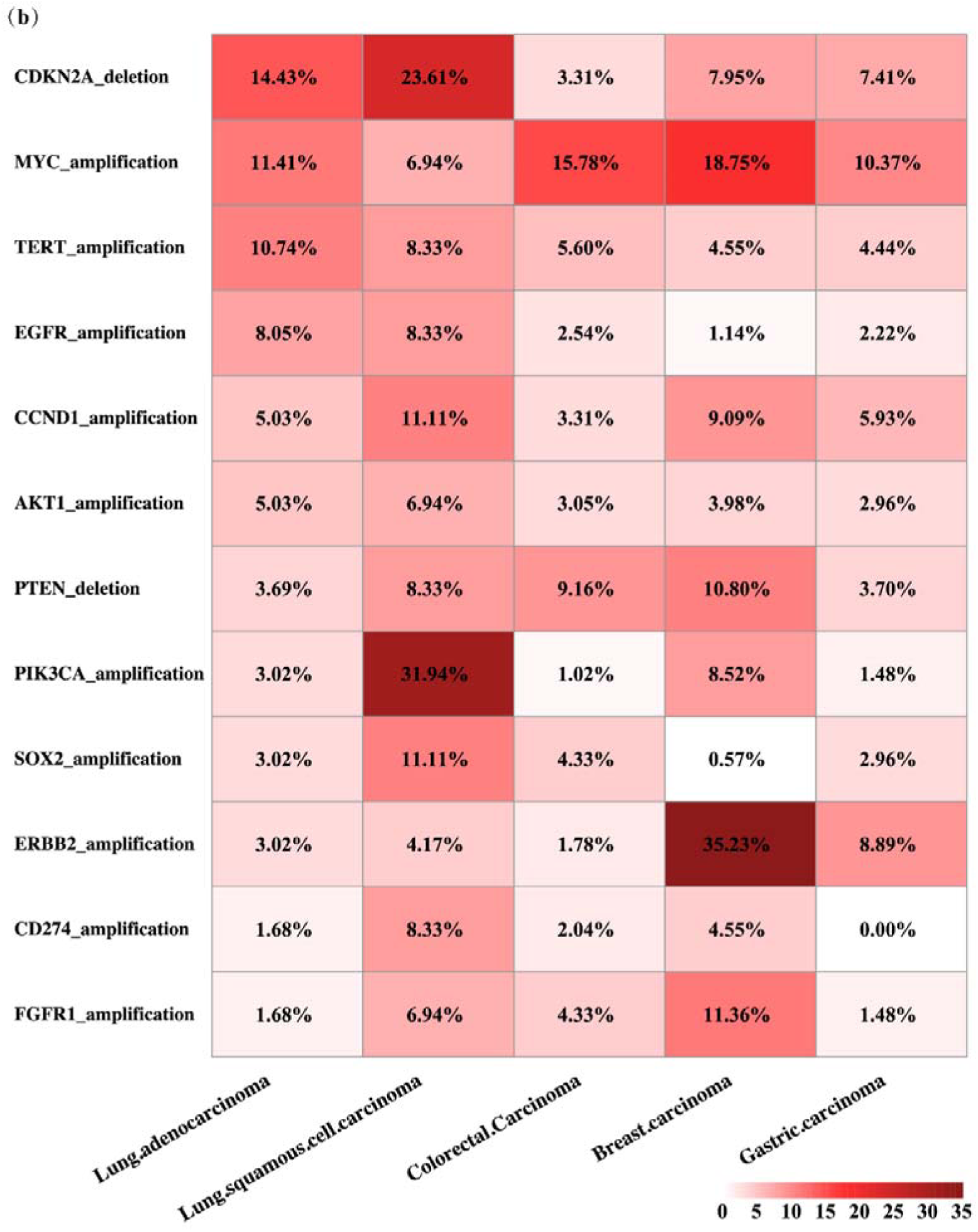
Frequency of somatic gene alterations of Chinese cancer patients. Genes with alteration frequency ≥ 1% are displayed. Bars indicate the percent of cases with distinct genomic alterations of each tumor type. (a) Single nucleotide variants (SNV). (b) Copy number variation (CNV).

### Analysis of the CWES results of Chinese cancer patients

According to data obtained from 953 CWES reports, 88.56% patients had at least one potentially actionable mutation and could be prescribed clinical treatment using precision medicine, based on the molecular mechanism detected in a tumor. This result was consistent with prior research, which reported values of 60.9% and 94.99%, respectively (Kou et al. 2017; Signorovitch et al. 2017). About 80% of patients were recommended for treatment with at least one FDA approved target/immunotherapy drug. About 40% of patients had at least 1 of the 4 potential immunotherapy predictive biomarkers (high TMB, MMR-deficient, high MSI and PD-L1 amplification). About 20% patients were recommended for approved indications of identified targeted drugs. Overall, CWES reports provided clinical relevant treatment recommendations for most of the patients (*Figure 4*).

**Figure 4.**
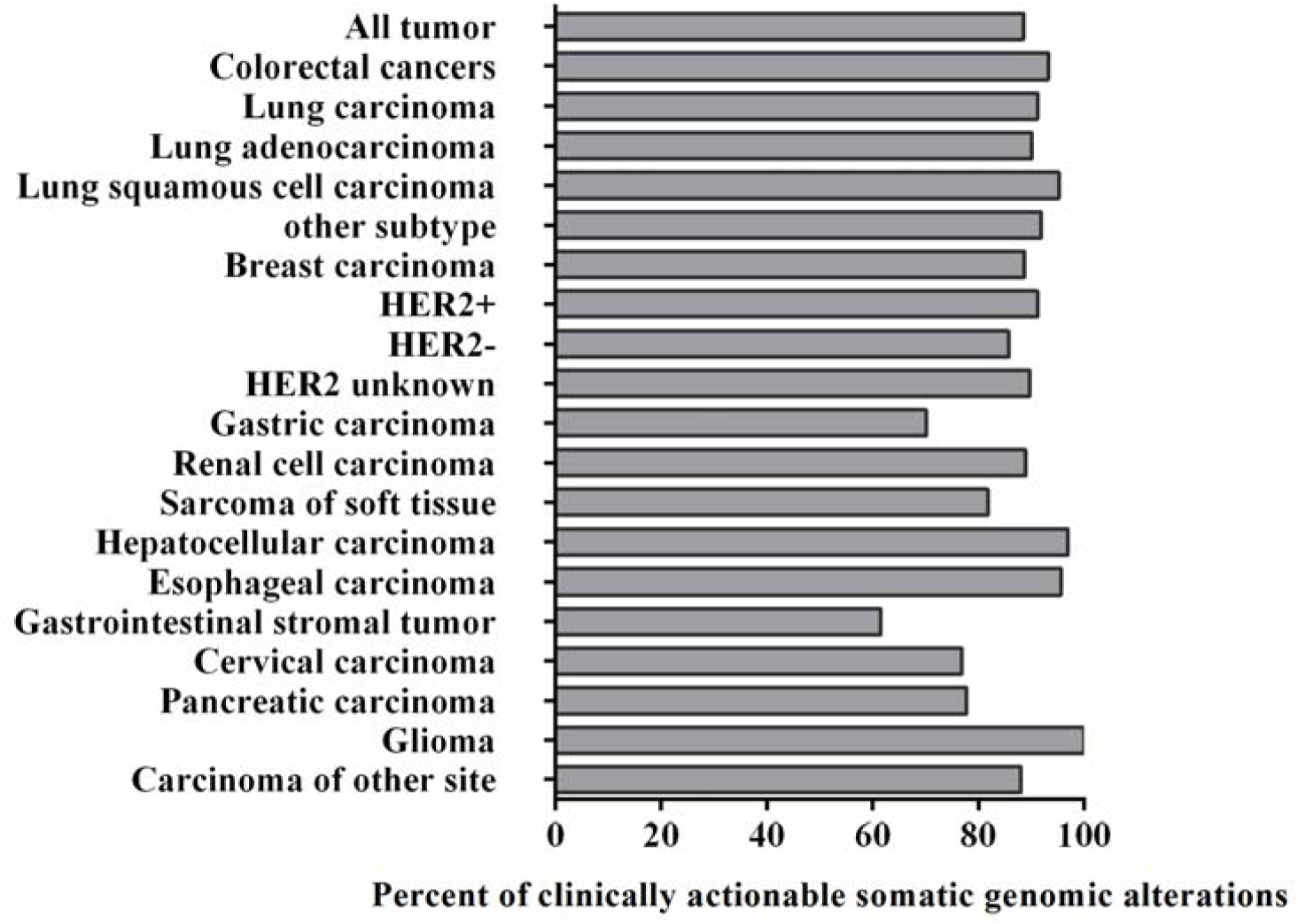
Clinically actionable somatic genomic alterations of Chinese cancer patients. Of these, 88.56% patients had at least one potentially actionable mutation and could be prescribed clinical treatment using precision medicine, based on the molecular mechanism detected in a tumor.

### Assessment of the clinical benefits of CWES

A total of 22 stage IV patients formed the study cohort, and their characteristics are described in *Table 1*. About 45% of the total were lung cancer patients (n = 10), and 27% had colon cancer (n = 6). Other types were breast, pancreatic, renal, thymus, fibroblastoma, mesothelioma, and hemangioendothelioma cancers.

**Table 1.**
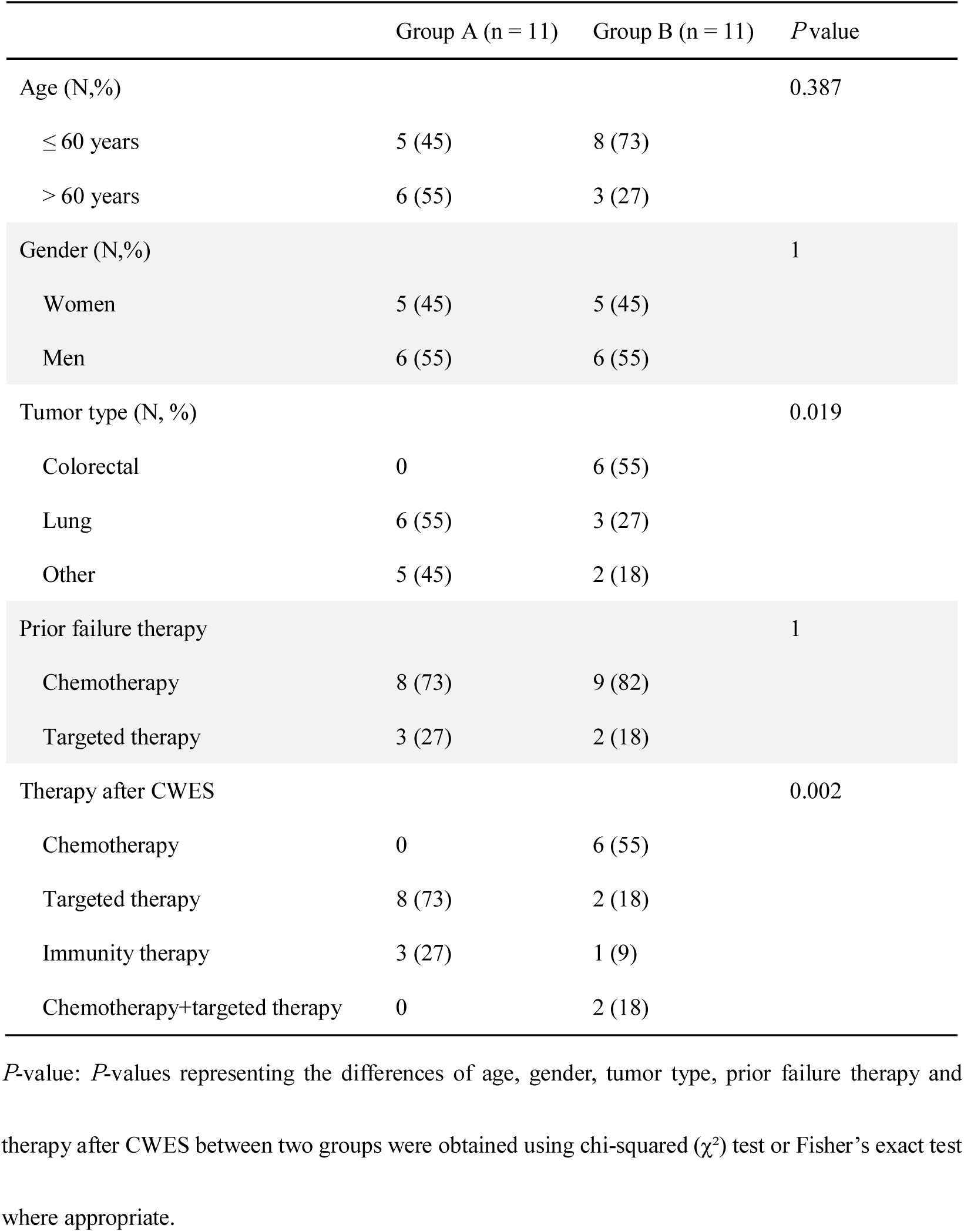
Demographic and clinical characteristics of Chinese cancer patients.

|  | Group A (n = 11) | Group B (n = 11) | *P* value |
| --- | --- | --- | --- |
| Age (N,%) |  |  | 0.387 |
| ≤ 60 years | 5 (45) | 8 (73) |  |
| > 60 years | 6 (55) | 3 (27) |  |
| Gender (N,%) |  |  | 1 |
| Women | 5 (45) | 5 (45) |  |
| Men | 6 (55) | 6 (55) |  |
| Tumor type (N, %) |  |  | 0.019 |
| Colorectal | 0 | 6 (55) |  |
| Lung | 6 (55) | 3 (27) |  |
| Other | 5 (45) | 2 (18) |  |
| Prior failure therapy |  |  | 1 |
| Chemotherapy | 8 (73) | 9 (82) |  |
| Targeted therapy | 3 (27) | 2 (18) |  |
| Therapy after CWES |  |  | 0.002 |
| Chemotherapy | 0 | 6 (55) |  |
| Targeted therapy | 8 (73) | 2 (18) |  |
| Immunity therapy | 3 (27) | 1 (9) |  |
| Chemotherapy+targeted therapy | 0 | 2 (18) |  |

*P*-value: *P*-values representing the differences of age, gender, tumor type, prior failure therapy and therapy after CWES between two groups were obtained using chi-squared ( $\chi^2$ ) test or Fisher's exact test where appropriate.

Patients who accepted treatment following the CWES reports were defined as group A, containing 11 patients; 11 patients did not follow the CWES suggestions (Group B). The types of therapies before CWES were similar in the two groups (8/11 chemotherapy, 3/11 targeted therapy in group A; 9/11 chemotherapy, 2/11 targeted therapy in group B). According to information at the last follow-up of each patient, in the A group, 11 (100%) patients received all precision therapy after CWES tests which included 8 patients for targeted therapy and 3 patients for immune-therapy, while in the B group, 6 patients still received chemotherapy, and only 2 patients accepted targeted therapy and one patient received immune-therapy (*Table 1*). We also analyzed statistical differences in demographic and clinical characteristics between the 2 groups, the tumor type and the next round of treatment differed significantly; *P*-values were 0.019 and 0.002, respectively. CWES affected the next round of treatment.

In group A, 1 patient achieved a complete response, 4 patients’ tumors shrunk, 1 patient achieved stable disease, and 4 patients progressed at the median time of 5 months. One patient stopped the treatment because of adverse reactions, even though the treatment was effective. Overall, 5 (45%) patients failed this round of therapy. In group B, 1 patient achieved stable disease and 10 patients progressed at a median time of 3 months, with a failure rate 91% for the treatment, lower than in group A (*Table 2*). PFS of group A had a median time of 12 months, but the median time of group B was only 4 months (*P* = 0.0016) (*Figure 5*). The longest PFS in Group B was 9 months, while almost half of Group A had a PFS beyond 9 months. In Group B, only 1 patient used PD-1 blockade pembrolizumab without genomic support for only 1-month of disease control. In Group A, 3 patients used pembrolizumab with genomic support, 2 with disease control for 12 months, and 1 with progression after 7 months. These results indicated that CWES could provide very effective treatment advice, and advanced cancer patients in China can benefit from CWES.

**Table 2.** Tumor Response

| Response | Group A (n=11) | Group B (n=11) |
| --- | --- | --- |
| Complete response (N, %) | 1 (9%) | 0 |
| Tumor shrinkage (N, %) | 4 (36%) | 0 |
| Stable disease (N, %) | 1 (9%) | 1 (9%) |
| Disease progression (N, %) | 4 (36%) | 10 (91%) |
| Adverse events (N, %) | 1 (9%) | 0 |

**Figure 5.**
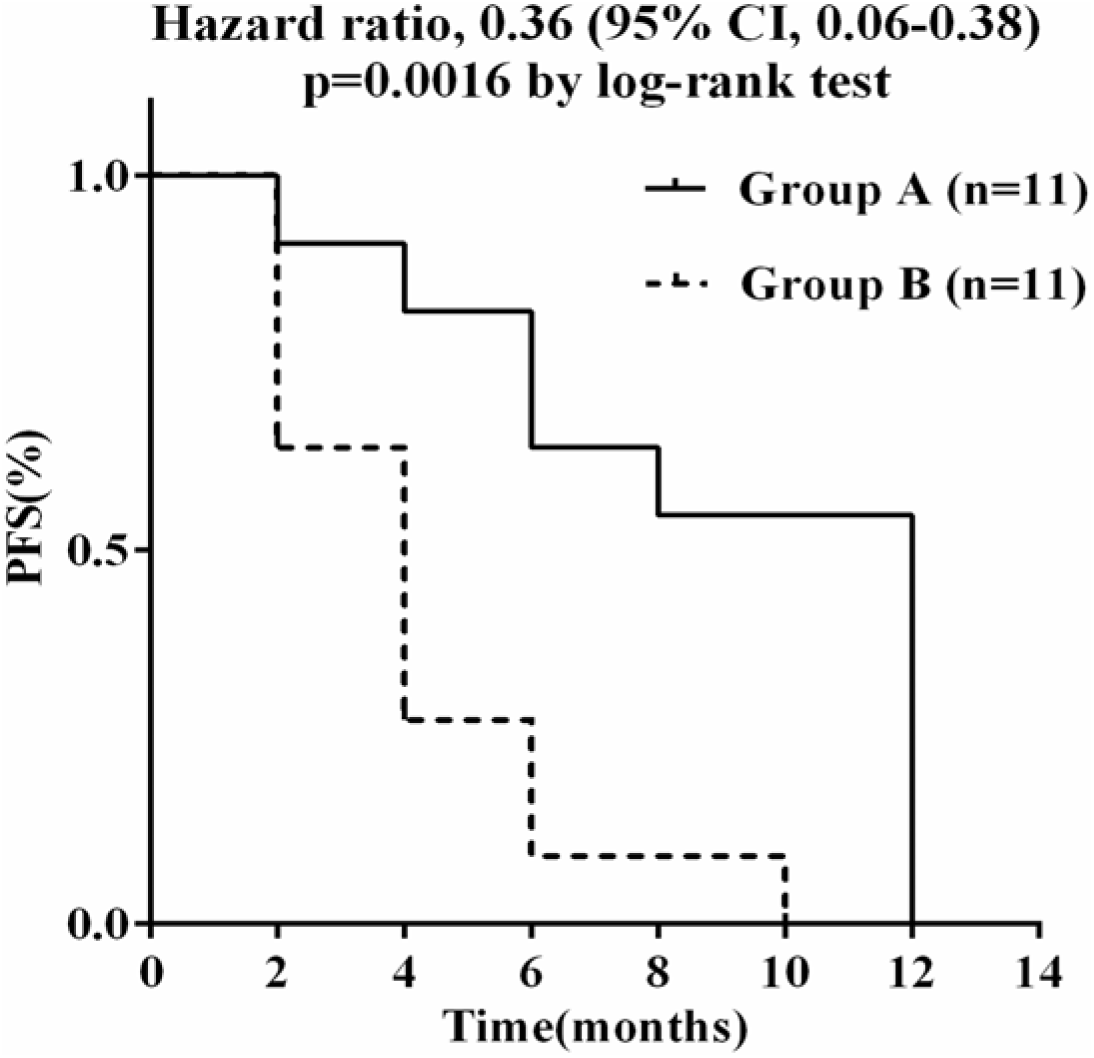
Kaplan-Meier curves for PFS. The median PFS of group A and group B are 12 and 4 months respectively.

## Discussion

A number of studies have considered whether the information obtained by high-throughput genomics from cancer patients could improve their outcomes. The SHIVA randomized trial revealed PFS was not significantly improved after precision medicine compared with the standard-of-care arm (Le et al. 2015). Perhaps this trial before 2015 lacked the experience acquired in previous precision medicine investigations, and did not use updated targeted drugs and immunotherapy. There are other reports that suggest that integrative high-throughput genomics and information of clinical therapy have positive effects for cancer patients. These studies originated in American, France and Japan, and included adults, young adults and children; most of the patients had hard-to-treat cancers or very advanced cancers (Kou et al. 2017; Massard et al. 2017; Mody et al. 2015a; Signorovitch et al. 2017). To the best of our knowledge, this is the first study assessing the clinical benefits of CWES strategy in China.

In the present study, we enrolled more than 1,000 Chinese patients with distinct types of cancer. We found differences in genomic profiling between the Chinese and TCGA. In addition to TP53, the frequency of mutated genes in gastric cancer patients in China was generally rather low, suggesting that the pathogenesis of gastric cancer is complex and heterogeneous. More importantly, we believe that patients with LUAD in China will derive a higher rate of benefit from EGFR TKI treatment, as the frequency of EGFR mutations in Chinese LUAD patients is much higher than that of TCGA. These genomic profiling data will provide information for the development of Chinese precision oncology.

Approximately 88.6% of patients had at least one potentially actionable mutation, and about 80% patients were matched with at least one targeted/immuno-therapy in our study. Our study was consistent with previous research (Kou et al. 2017; Signorovitch et al. 2017). But the percentage of actionable alterations was higher than in the MSK-IMPACT test, where 36.7% of patients were shown to harbor at least one actionable alteration (Zehir et al. 2017). Relatively speaking, colon, lung, breast and gastrointestinal stromal tumors have more opportunities to receive targeted therapies and immunotherapy than other forms of cancer. First, in our tests, these types accounted for 78%, but only 44% in the MSK-IMPACT test. Second, except for information from FDA and off-label treatment, we collected clinical information from all investigational clinical trials, but the MSK-IMPACT test only contained compelling clinical evidence.

This study may bring hope to Chinese patients with advanced cancer. We found CWES can significantly affect the choice of follow-up treatment for Chinese cancer patients (*P* = 0.002). More importantly, we also found that cancer patients can benefit from CWES guiding based on follow-up data. 4 patients progressed, 6 (55%) patients obtained disease control with following CWES reports.

But only 1 patient achieved disease control without following CWES guidelines, and others all progressed. PFS was significantly longer when following the CWES group compared to the non-followed patients (median PFS 12 vs 4 months, *p* = 0.0016). Due to all colon cancer patients without following CWES guidelines, we left colon out and recalculated the PFS. PFS of group A had a median time of 12 months, and the median time of group B was only 2 months (*P* = 0.011) (*Supplementary Figure 1*). It also differed significantly. These results are consistent with previous research reports (Massard et al. 2017; Mody et al. 2015b; Schwaederle et al. 2016; Signorovitch et al. 2017). In a study of 102 children and young adults with relapsed, refractory or rare cancer, 10% of patients accepted personalized clinical interventions, leading to remissions of between 8 and 16 months or which produced sustained complete clinical remission of 6 to 21 months (Mody et al. 2015b). In the MOSCATO 01 clinical trial with a cohort of 1,035 cancer patients, 24% of patients were treated with a targeted therapy matched to a genomic alteration. It is noteworthy that 1% of patients had complete responses, and 52% patients reached stable disease (Massard et al. 2017). Although the present study and others showed different degrees of effective treatment, they strongly indicated that information integrating an individual genomic sequencing and treatment strategy will improve the clinical outcomes in a subset of patients.

Cancer is a disease of the genome (Garraway et al. 2013; Rubin 2015). Each patient has his or her own unique pathological genotype and it is difficult to make an accurate diagnosis without thorough gene sequencing. For example, some oncogenes are main driver genes in some types of cancers, but these oncogenes are rare in other cancers. Fusions of EML4-ALK are seen in 3–5% of lung cancer patients, but they are extremely rare in papillary thyroid cancer (Demeure et al. 2014). And some oncogenes are less frequently expressed, such as the fusions of neurotrophic receptor tyrosine kinases (NTRK) that only have an approximate 0.5% probability of being expressed in diverse solid tumors and hematological malignancy, but these are also fatal for many cancer patients. The patients who received targeted NTRK-fusion treatment had obviously improved response rates (Vaishnavi et al. 2015). Most of the frequent cancers present with a large number of very rare but targetable genomic alterations (Lawrence et al. 2014). In addition, multiple accumulated genomic cancer knowledge produces an avalanche of targeted and immune therapies to be tested in clinical trials, and a rapidly growing list of cancer drugs has been approved that match specific genetic alterations. CWES can provide this information. These cases indicate CWES is important for every patient, taking into account individual differences.

Our study has a number of limitations. It did not provide a higher level of evidence for our hypothesis. Randomized trials and real world studies will be required to validate our hypothesis and to quantify the magnitude of benefit. There are several ways to improve further the efficacy of our strategy. RNA sequencing could define pathway activation and target gene expression levels. Detection of circulating tumor DNAs (ctDNAs) of blood provides a non-invasive and easily accessible way for monitoring cancer progression. Therefore, the combination of target gene panel, WES, RNA sequencing plus detection blood ctDNAs will be a powerful tool for cancer treatment.

## Conclusions

The current study not only revealed differences in genomic profiling between Chinese cancer and TCGA, it also showed that CWES can provide clinically relevant treatment recommendations for most Chinese cancer patients. Furthermore, our preliminary observations suggest that advanced Chinese cancer patients will benefit from CWES.

## Author contributions

QX, SJ and YB contributed to the design and conduct of the study. SJ and YB were responsible for the collection and analysis of clinical data. CD, XC, and XX performed the experiments. GW and JW were responsible for bioinformatics and statistical analyses. BW and WS was responsible for writing the manuscript. All authors approved the final version of the manuscript.

---

**Supplementary Figure 1.**
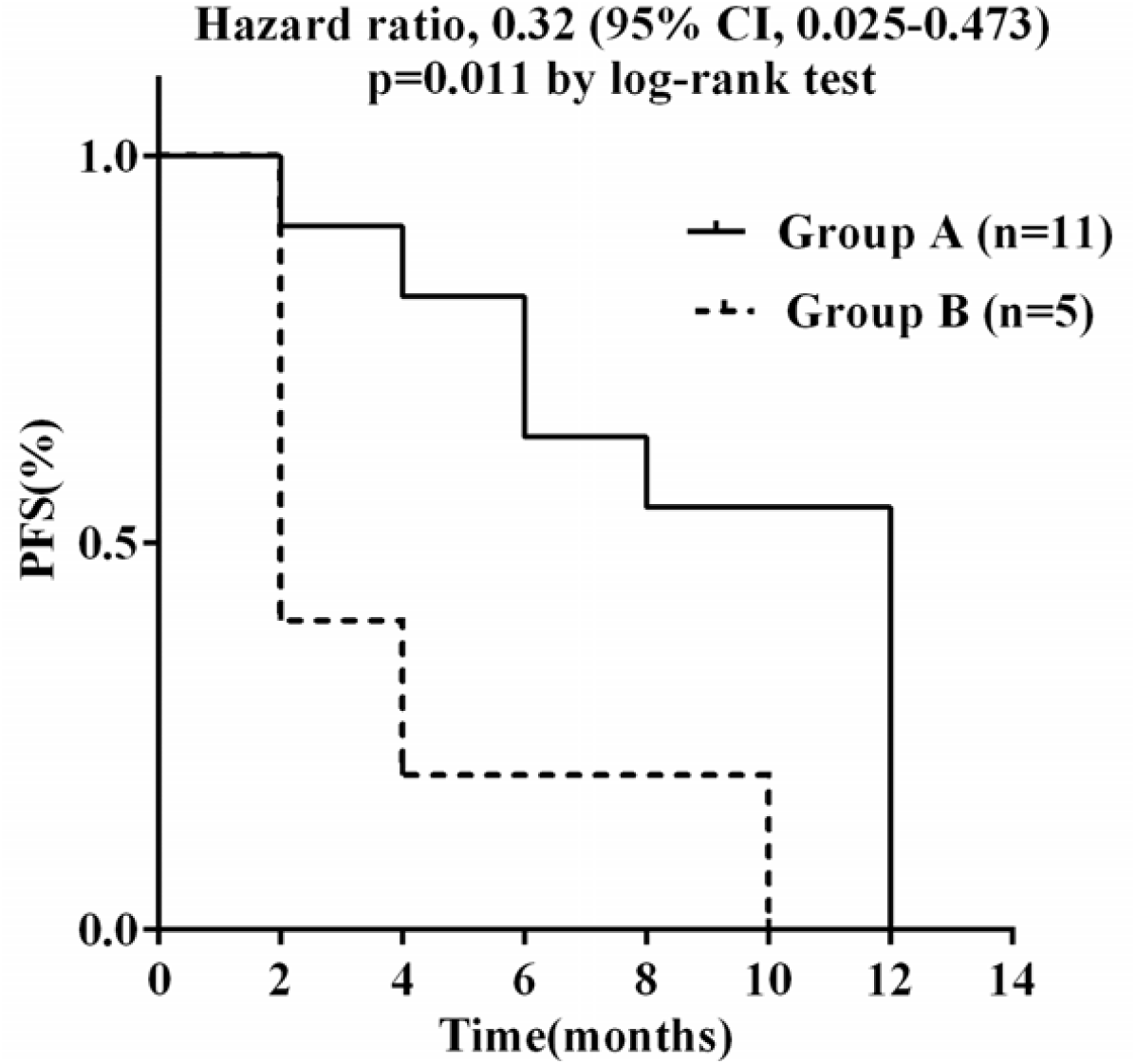
Kaplan-Meier curves for PFS without colon cancer. The median PFS of group A and group B are 12 and 2 months respectively.

